# Network analysis after epigenome wide methylation study revealed JUP as a regulator of co-methylated risk-module for T2DM

**DOI:** 10.1101/2021.07.01.450660

**Authors:** Anil K Giri, Gauri Prasad, Vaisak Parekkat, Donaka Rajashekar, Nikhil Tandon, Dwaipayan Bharadwaj

## Abstract

Controlling the global Type 2 diabetes mellitus (T2DM) pandemic requires a better understanding of its risk factors across different populations, and needs markers that can precisely predict the individual risk to the disease. DNA methylation due to environmental factors is a known mechanism for conferring risk to T2DM. To identify such methylation signatures and associated risk to the disease, we performed an epigenome-wide-association study (EWAS) in 844 individuals of Indo-European origin. Within the Indian population, we identified and validated 6 novel differentially methylated CpG sites in *PDCD6IP*, *MIR1287*, *5S_rRNA*, *HDAC9*, *KCNK16*, and *RTN1* genes associated with T2DM risk at the epigenome-wide-significance-level (p<1.2×10^-7^). Further, we also replicated the association of 3 known CpG sites in *TXNIP*, *SREBF1* and *CPT1A* in the Indian population. Using methylation-based-network analysis, we identified 4 co-methylated modules, which we named as turquoise, yellow, brown, and blue, among differentially methylated CpG sites in discovery phase samples. We observed that methylation of the brown module with 28 CpG sites, associated with T2DM risk factors (e.g., BMI, insulin, C-peptide). Upon further analysis, we noted that these methylation signatures at 14 of the brown module’s CpG can be used as marker to segregate T2DM patients with good glycemic control (e.g., low HbA1c) but poor lipid profile (low HDL and high TG) from the other patients. Additionally, we discovered that rs6503650 in the *JUP* gene regulates methylation at all the 14 hub CpG sites of the brown modules as methQTL. Our network-assisted epigenome-wide association study is the first to systematically explore DNA methylations conferring risks to T2DM. In addition, the study shows the potential use of identified risk CpG sites for patient segregation with different clinical outcomes. These findings can be useful for better stratification of patient in order to improve the clinical management and treatment outcomes.

## Introduction

Type 2 diabetes mellitus (T2DM) is a causal factor for several micro-(e.g., retinopathy, nephropathy, neuropathy) and macro-vascular related complications (e.g., cardiovascular disorders) with nearly 4 million T2DM related fatalities each year (1, 2). T2DM poses a major global health challenge for the modern health system as one-tenth of every dollar spent on global healthcare is on treating T2DM related complications (1). The devastating effect of T2DM on global health highlights the need to understand its risk factors in order to design better prevention plans and treatment strategies.

Existing evidence from genetic epidemiological studies and randomized control trials designed to intervene with T2DM risks suggest its multifactorial origin with significant contribution from genetic, environmental, and behavioral factors (3,4). Although, many of these causal factors are individual-specific (e.g., smoking, exercise) (5, 6,7,8), several others are population-specific (e.g., nature and content of diets, genetic component) (4, 9) as well as environment-specific (e.g., gut microbiome, sedentary lifestyle) (10,11) that significantly differ across different ethnic populations. These behavioral and environmental factors contribute to diabetes by controlling the gene expression through epigenetic mechanisms (e.g., DNA methylation, histone modification, chromatin remodeling) (12,13). For instance, exercise can cause hypermethylation of the DNA promoter in lipogenesis-associated HDAC4 and NCOR2 genes, leading to their decreased expression in humans (14). Further, DNA methylation affects expression of genes related to T2DM pathophysiology in pancreatic β-cell (15), adipose tissue (16), and skeletal muscle (17,18). Thus, identification of mechanisms showing how DNA methylation variations present in individuals across different ethnicity confer risk to T2DM is important to better understand the disease biology (19).

Previous epigenome-wide association (EWA) and candidate-based studies have identified more than 60 differentially methylated CpG sites in more than 40 genes (e.g., TXNIP) as risk loci for T2DM (http://ewascatalog.org/?trait=type%20ii%20diabetes). However, these individual CpG sites have a small influence over T2DM risk as observed from their small effect size and minor methylation differences at the loci between patients and healthy controls. The small effect size of the CpG sites further indicates that a coordinated methylation or demethylation at multiple CpG sites in a specific region (e.g., promoter, gene body) of a gene or group of genes is required to bring the necessary changes in disease-related gene expression (20). Hence, it is crucial to study the group behavior (co-methylation) of identified CpG sites to decipher the coordination among respective genes affecting the disease process. However, despite the growing literature describing the role of DNA methylation in T2DM biology, the co-methylation behavior of the identified CpG sites, and their role in T2DM risk is largely unknown (19).

To fill the gap in knowledge, the current study firstly identifies differentially methylated CpG sites between healthy controls and T2DM patients in 844 subjects of Indo-European origin from Indian population. Our results replicated 3 earlier known signals in the Indian population and identified 6 novel differentially methylated CpG sites in the T2DM patients that also associate with other T2DM-risk factors (e.g., BMI, lipids). Secondly, our study utilized the Weighted gene expression network analysis (WGCNA) (21), a commonly used systematic algorithm, to identify the co-expressed group of genes which significantly affects the disease risk from a dataset consisting of thousands of genes. WGCNA has been used to identify genes, microRNAs (miRNAs), and long noncoding RNAs (lncRNAs) significantly correlated with progression, prognosis, metastasis, and recurrence in multiple diseases (22,23,24). With the help of WGCNA, we identified 4 co-methylated modules among the differentially methylated CpG sites. We further explored the mechanism for regulation of methylation in CpG sites of identified modules using methylation-quantitative trait loci (methQTL) analysis. Consequently, we identified a novel co-methylated module consisting of 28 differentially methylated CpG sites with the *JUP* gene as the central node. All the hub genes in the module were regulated by a methQTL (rs6503650) in the *JUP* gene.

## Methods

### Study population and measurements of clinical parameters

The ethnicity of every study participant (Indian) is of Indo-European origin and they are member of the INdian DIabetes COnsortium (INDICO) (25,26). T2DM patients have been recruited from Diabetes Clinics from the Department of Endocrinology, Metabolism & Diabetes, All India Institute of Medical Sciences (AIIMS). We defined T2DM according to the World Health Organization criteria, as a condition with any of the following characteristics: (i) fasting glucose level of ≥126 mg/dl (7.0 mmol/L), (ii) 2-hour postprandial glucose ≥200 mg/dl, (iii) glycosylated HbA1C level > 6.5%. Diseased subjects with an age of onset below 30 years were also excluded to avoid mix-up with type 1 diabetes.

The controls have been recruited by organizing health awareness camps in the community (27,28) around Delhi and nearby areas and were defined as (i) >40 years of age; (ii) HbA1c level <6.0%; (iii) fasting plasma glucose level <110 mg/dL (6.1 mmol/L); (iv) no family history of diabetes in first-degree relatives; and (v) urban dwellers. Informed written consent was obtained from all the participants of the study. The ethnicity of the subject was defined primarily based on the geographical location of the recruitment along with their birthplace and place of origin of both the parents. Pregnant women and individuals of other ethnicities were excluded from the study. The characteristics of the study participants have been shown in Supplementary Table 1. All study participants have been well characterized for their anthropometrical (e.g., height, weight, waist circumference, and hip circumference), and biochemical measures (e.g., fasting glucose, glycosylated hemoglobin (HbA1C), insulin, C-peptide, total cholesterol, HDL, LDL, triglycerides, C-reactive protein (CRP), as mentioned previously (26, 29,30). Questionnaires assessing cigarette smoking status (coded as current smoker/ex-smoker/never), alcohol consumption status (coded as regular/sporadic/never), and level of leisure-time physical activities (coded as regular/sporadic/never) were recorded. The study was approved by the ethics committees of the participating institutes and carried out in accordance with the principles of the Helsinki Declaration.

### Discovery phase DNA methylation assay and quality control analysis

We performed the Infinium assay using HumanMethylation450 BeadChip for 524 (260 T2DM+264 controls) Indian individuals of Indo-European origin. The genomic DNA required for the assay was extracted from peripheral blood leukocytes using the salting-out process as mentioned previously (31). Bisulfite conversion of 700 ng genomic DNA was performed using EZ-96 DNA Methylation Kit (Deep Well Format) (Zymo Research; Irvine, CA, USA) in accordance with the given kit instructions (www.zymoresearch.com). The Infinium assay for genome-wide DNA methylation using HumanMethylation450 BeadChip was performed as per the manufacturer’s instruction. Briefly, bisulfite-converted DNA was sequentially alkali denatured, neutralized, isothermally amplified by incubating at 37°C for 22 hours, end-point enzymatically fragmented, purified, and hybridized on an Infinium^®^ HumanMethlyation450 BeadChip at 48°C for 18 hours, as described previously (31,32). The BeadChip was washed to remove any free or non-specifically hybridized DNA; the hybridized probes were single-base extended and was stained with multiple layers of fluorescence. Finally, BeadChip was coated and scanned using Illumina^®^ iScan system. The image files were analyzed using the minfi package in R (Supplementary Figure 1) (33). The analysis involves background subtraction, fixing the outlier methylation values using “fixMethOutliers” option in the minfi package. We removed samples which could not pass the quality control (QC) measures. We excluded 2 samples with a call rate <95% for probes, 4 poor-quality samples having at least 1% of CpG sites with detection *p*-value less than 0.01, 1 sample with bisulfite failure (intensity lesser than 3 SD of the mean value for C1, C2, C3, and C4 probes present in the chip) and 1 sample with possible gender mismatch based on evaluation of selected CpG sites on the Y chromosome, leaving a total of 504 samples available for further analysis (Supplementary Figure 1). At the CpG sites level, we removed 457 CpG sites that had a bead count <3 in 5% of samples, 831 CpG sites with average detection p-value of >0.01 in 1% of samples and 570 CpG sites with call rate <98%. We also removed 30903 cross-hybridization probes present in the supplementary file of Chen et al [34] and 15781 CpG sites containing SNP on the probe. In addition, we removed 10277 CpG sites present in sex chromosomes. The analysis plan details along with various quality control steps, number of samples and CpG sites removed, have been shown in Supplementary Figure 1.

The data was normalized using the beta-mixture quantile normalization (BMIQ) model to remove the technical variability inherent in two different types of probes present in HumanMethylation450 BeadChip (35). We estimated different WBC cell compositions using the Housman Algorithm based on methylation level of 500 CpG and used linear regression to regress out the effect of age, sex, smoking status, alcohol consumption, diet, bisulfite conversion efficiency, and cell compositions over methylation level of each probe (36). Wilcoxon test was used to compare the mean methylation difference between T2DM subjects and healthy controls for 426693 CpG sites in 504 subjects (254 T2DM subjects and 250 controls) using IMA packages in R. A p-value of < 1.2 × 10^−7^ (0.05/426693) was considered significant after Bonferroni correction for the number of CpGs. The inflation factor for the association was 1.67 that is apparently high but genomic inflation is often discussed as endemic to epigenome-wide association studies (EWAS) (37, 38).

### Selection of CpG sites for replication

We validated the top identified signals using the EpiTYPER (39) assay in 320 individuals (157 T2DM + 163 control). We selected a total of 17 CpG sites for validation, which included the top 4 differentially methylated CpG sites based on p-value and 1 site with maximum β-differences between case and control groups as significant CpG sites with an assumption that a larger difference could also influence the disease (Supplementary Table 2, Supplementary Figure 2).

We also used data-driven approaches for CpG selection for the validation phase. To identify T2DM associated CpG sites with the effect on other T2DM related traits, the association of 535 CpG sites was sought in controls, with extreme distribution for T2DM related traits such as glycaemic parameters, obesity-related parameters, lipid-related parameters, and insulin-related parameters. We compared the methylation level of the 535 identified CpG sites in subjects with a measurement below 10^th^ percentile and above 90^th^ percentile for extreme level for T2DM risk related phenotypes. This approach was supplemented with the role of corresponding genes in relation to respective phenotypes in literature. Based on this approach, we selected 17 signals for the validation phase using EpiTYPER assay (39).

### Validation of significant signals from discovery phase in an independent set of samples using the EpiTYPER assay

For EpiTYPER assay, bisulfite-converted primers corresponding to 600 base pair passing the selected CpG sites were designed using EpiDesigner software (http://www.epidesigner.com/) (39). Bisulfite converted DNA (700ng) was performed using Zymo EZ-96 methylation kit by following the manufacture’s protocol. PCR reactions were performed in duplicates using bisulfite DNA and Agena PCR chemicals, followed by *in-vitro* transcription and uracil-based cleavage using RNAse A. The cleaved products were assessed using “time of flight” mass spectroscopy and methylation levels were quantified. The data was extracted and subjected for various QC steps using “MAssArray” package in R. We removed CpG sites if they have called <80% of subjects among total subjects. CpG sites having overlapping masses with other CpG sites were also excluded. Samples showing >20% of CpG sites as missing, were also discarded. Next, an average methylation value from PCR duplicates was estimated and used for further downstream analysis. We considered a p-value less than 2.94×10^-3^ (0.05/17) for replication of signals in the 2^nd^ phase considering multiple comparisons for 17 tests.

#### Meta-analysis of the summary statistics from the discovery and validation phase

We meta-analysed the replicated signals using the *metacont* function in “meta” packages in R (40).

##### Construction of weighted gene co-methylation network of top signals

Since network-based analysis allows assessing combined effect of differentially methylated CpG sites over disease risk, we constructed a weighted undirected co-methylation network in 264 controls using the methylation data for 535 CpG sites differentially methylated (p<1.2×10^-7^) between T2DM and healthy subjects using the WGCNA package in R (21). Within the network, nodes represented significant CpG sites and edges between nodes represented correlation between the CpG sites. We used a soft threshold power of 6, minimum module detection size of 10, fixed height cut-off to 0.25, and biweight mid-correlation as a measure of similarity and all other default settings for network construction. We defined modules in the network using the dynamic tree-cutting algorithm on the dendrogram, obtained from the hierarchical clustering option as provided in WGCNA. Furthermore, the connection strength between two nodes (CpG sites) of the module was estimated as the overlap in connection pattern between two nodes. Briefly, less connection strength denotes a poor correlation between module members. The gene connectivity is calculated by taking the sum of connection strengths with other genes. Module members with connectivity >0.80 were defined as hub CpG sites. Module members with connectivity <0.1 were removed from the modules to visualize the module structure clearly.

Lastly, we validated the preservation of modules in 254 T2DM subjects (Figure 2b) and another independent dataset of 55 individuals (28 individuals of Northern European ethnicity living in the USA for several generations and 27 migrant Indians to the USA who migrated for at least five years) (data not shown) (41) using the “modulePreservation” command in WGCNA (21). WGCNA calculates Z-summary, which is the measure of preservation of average of density measures (Zdensity) and connectivity measures (Zconnectivity) between the test and reference network. The higher the Z-summary score the higher the preservation and median preservation rank. Visualization of module structure and topology of the brown module was done using Cytoscape (42).

**Figure 1:**
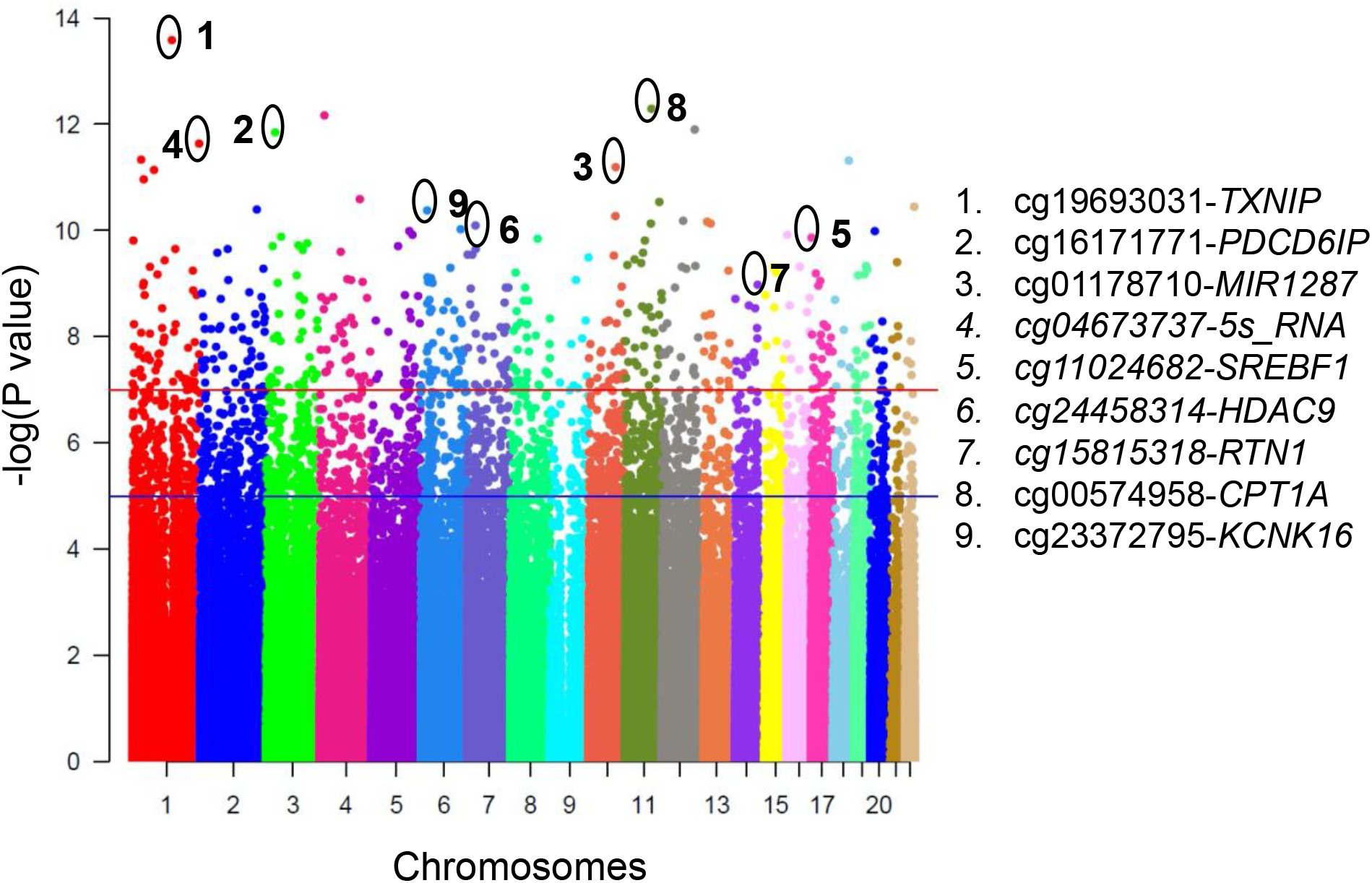
Manhattan plot showing the distribution of p-values for analyzed markers across different chromosomes. Manhattan plot for CpG sites differentially methylated between T2DM and healthy control in the discovery phase. The −log10 p-values for the association of CpG sites are plotted as a function of genomic position (National Center for Biotechnology Information Build 38). The p-values were determined using Wilcox Test in the discovery phase of analysis. Each chromosome (Chr) has been represented with a unique color.

**Figure 2:**
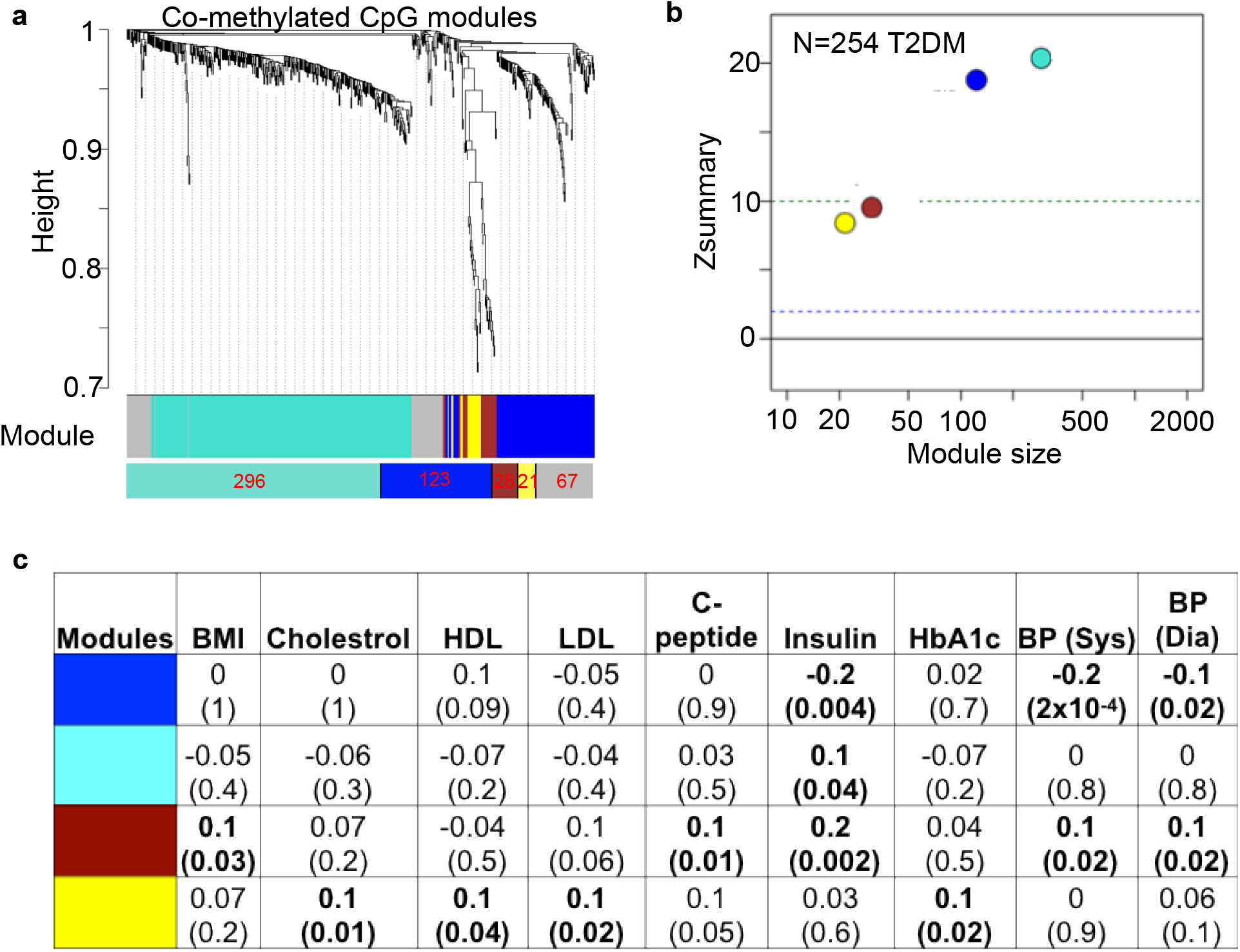
(a) Co-methylated modules among significant CpG sites in the discovery phase of EWAS. WGCNA identified 4 modules (represented by 4 colors-turquoise, blue, brown, yellow) CpG sites not assigned to any module have been shown in grey. (b) Preservation of identified modules in 254 T2DM cases has been shown in a scatter plot between Zsummary and module’s size. It indicates that the turquoise module has the highest level of Zsummary followed by blue, brown, and yellow modules, respectively. (c) Module–trait relationships. The rows represent the co-methylation module eigengene (ME) and its color. The columns represent clinically relevant traits in healthy individuals. The correlation has been shown as the upper number and P values have been shown as a lower number within each cell. The significant correlations have been marked in bold letters.

**Figure 3:**
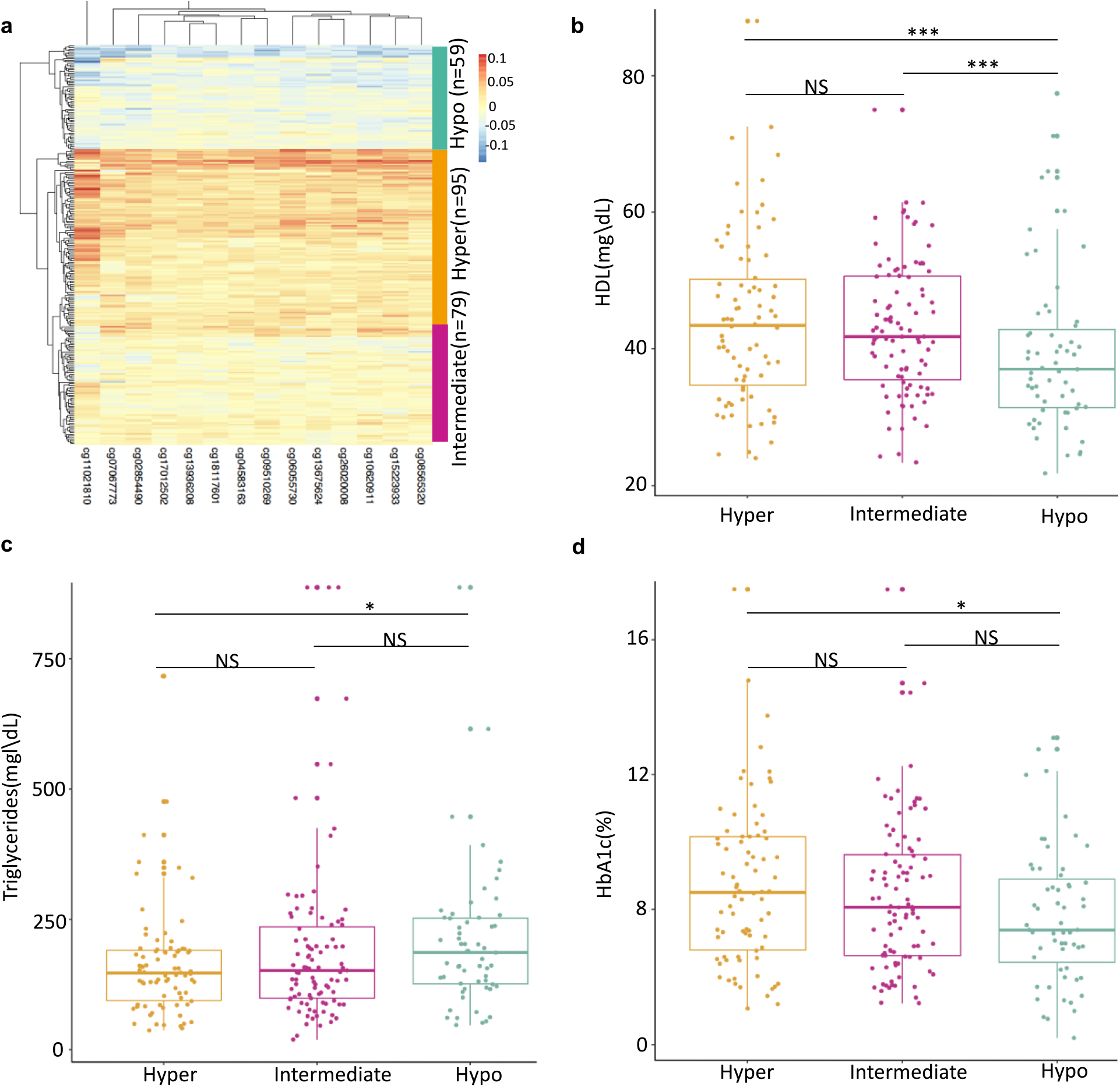
a. Heatmap showing clustering of T2DM patients using the methylation level of 14 hub CpG sites. The analysis revealed three subgroups of T2DM patients with low, intermediate, and high methylation at the hub CpG sites. Boxplot showing the difference in (b)HDL level, (b)triglycerides, and (c) HbA1c between 3 patients’ groups. The p-value has been calculated using the non-parametric Wilcox test.

#### Construction of module-trait relationships and identification of hub CpG sites associated with clinical modules

Pearson’s correlation coefficients and P-values between modules eigenvalues and T2DM-related traits were calculated using WGCNA packages and visualized in a module-trait table. The higher the Pearson Co-relation Coefficients between eigengenes and the clinical variable will reflect the stronger role of module methylation in relation to phenotype. A module that affects a larger number of risk factors could be interesting for further mining and functional studies. A correlation coefficient with a p-value <0.05 was considered significant.

#### Functional enrichment analysis of the CpGs associated genes from the key modules

We performed pathway analysis and gene enrichment analysis for identified modules using GENECODIS to get an insight about the possible mode of action of identified modules in the biology of T2DM (43).

### Power analysis

We calculated 100% power of the study using “pwr” packages in R. Power analysis was performed for 500 samples using a significance level of 1×10^-7^, average β-difference of 0.01 and, the average standard deviation for probes was 0.026.

## Results

### Epigenome-wide association study for T2DM in Indians identified 6 novel loci in Indian populations

The result showed differential methylation for 535 CpG sites at a p-value of 1.2×10^-07^ (0.05/426693). The list of differentially methylated CpG sites has been shown in Additional file 1. Distributions of p-values across different chromosomes have been shown in the Manhattan plot (Figure 1). We detected a CpG cg19693031 in *TXNIP* as the top signal in the Indian population. Replication of 17 selected CpG sites using EpiTYPER assay was performed in 320 independent samples (157 T2DM+163 controls) and data was analyzed using the “MassArray” package in R. We observed a strong correlation (Pearson correlation of 0.76 at a p-value of 2.69×10^-5^ for cg01178710 in *MIR1287* gene, median correlation of 0.63) between Illumina 450K technology and EpiTYPER assay measurements (Supplementary Figure 3).

Meta-analysis of summary statistics from both the phases revealed differential methylation of 9 CpG sites between healthy and T2DM patients at the epigenome wide significance level in Indian population (p-value of <1×10^-7^, Table 1) (44). We also replicated signals cg19693031 in *TXNIP* (β = −0.05, p-value = 2.16×10^-16^) and cg11024682 in *SREBF1* (β = 0.02, p-value = 3.02×10^-10^) and cg00574958 in *CPT1A* (β = −0.01, p-value = 1.43×10^-8^) in Indian population living in India (Table 1). We identified 6 novel signals viz cg16171771 in *PDCD61P*, cg01178710 in *MIR1287*, cg04673737 in *5s_rRNA*, cg24458314 in *HDAC9*, cg23372795 in *KCNK16*, and cg15815318 in *RTN1* as differentially methylated in case of T2DM in Indian population (Table 1).

**Table 1:**
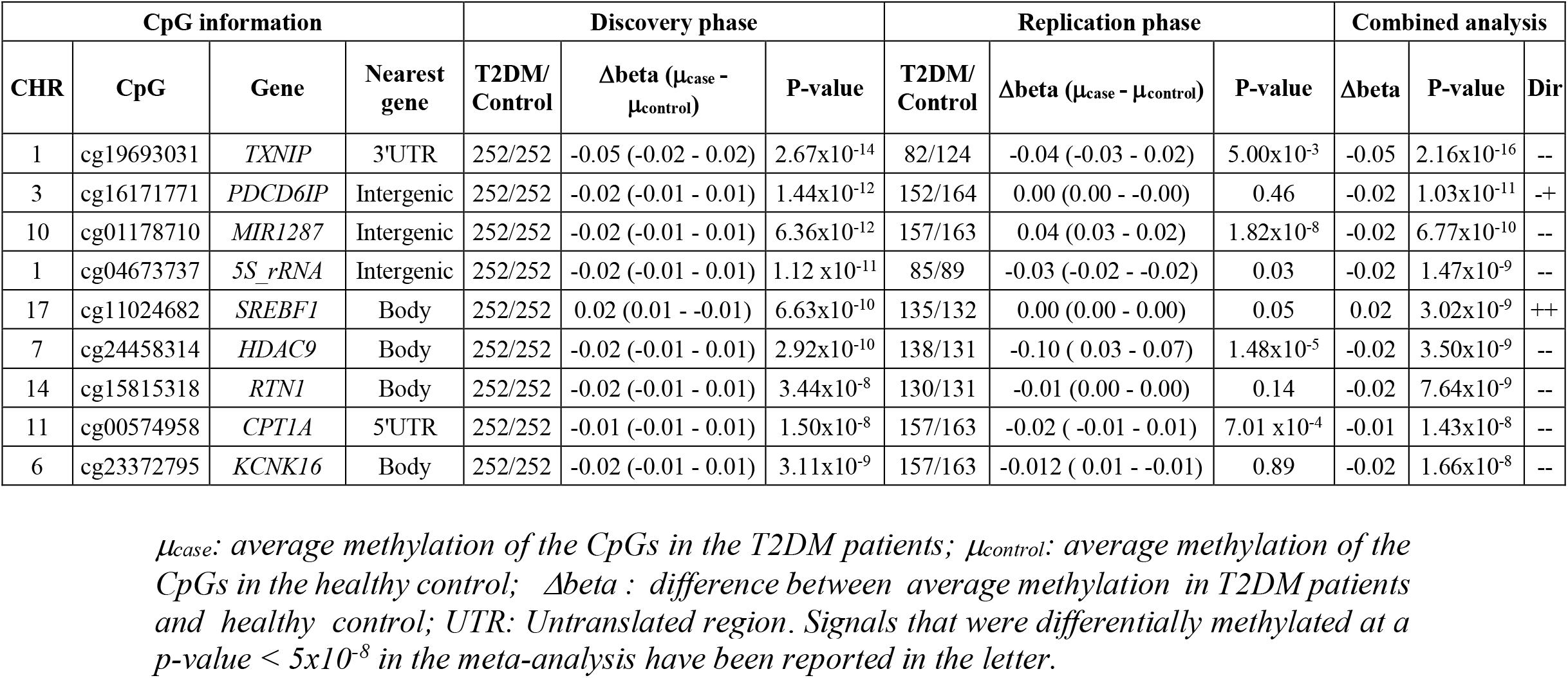
CpG sites showing differential methylation at epigenome-wide significance levels in the meta-analysis.

### Identified CpG sites associates with T2DM risk factors

To understand the possible action mechanism in the observed differential methylation over T2DM risk, we performed an association analysis of the identified signals with other T2DM risk factors (e.g., BMI, lipids, etc) in cases and control subject separately. The hypothesis being that DNA methylations at these significant loci alter the level of T2DM risk factors. The analysis revealed a significant association of identified CpG sites with fasting glucose, triglycerides, LDL, CRP, uric acid, creatinine, insulin, and obesity measures (e.g., BMI). For example, cg19693031 in *TXNIP* associated with fasting glucose, HbA1c, cholesterol, triglycerides, and LDL level in T2DM cases and WHR measurements in control subjects (Table 2). We observed an association of cg19693031 in *TXNIP* with hyperglycemia in Indian population similar to the Caucasian population from Spain (45). The association of identified signals with T2DM risk factors suggest a shared epigenetic contributor amongst them.

**Table 2:** Table showing association of methylation of identified CpG sites with other T2DM risk factors in healthy individuals.

|  |  | Controls |  | T2DM |  |
| --- | --- | --- | --- | --- | --- |
| CpG sites | Trait | Effect(SE) | P | Effect(SE) | P |
| cg19693031 (TXNIP) | Fasting glucose | -27.80(14.82) | 0.06 | -220.5 (59.26 ) | 2.49x10<sup>-4</sup> |
| cg19693031 (TXNIP) | HbA1c | -0.96(0.59) | 0.11 | -12.52 (2.01 ) | 1.92x10<sup>-09</sup> |
| cg19693031 (TXNIP) | Cholesterol | -81.28(51.42) | 0.12 | -244.67 (44.37 ) | 8.88x10<sup>-08</sup> |
| cg19693031 (TXNIP) | Triglycerides | -168.04(93.46) | 0.07 | -650.79 (124.52 ) | 3.66x10<sup>-07</sup> |
| cg19693031 (TXNIP) | LDL | -39.90(44.08) | 0.37 | -171.67 (34.99 ) | 1.74x10<sup>-07</sup> |
| cg19693031 (TXNIP) | WHR | -0.24(0.12) | 0.04 | 0.007 (0.05) | 0.89 |
| cg16171771 (PDCD6IP) | Triglycerides | -5.15(113.35) | 0.96 | -486.83 (215.12 ) | 0.02 |
| cg16171771 (PDCD6IP) | LDL | -106.77(52.78) | 0.04 | -42.36(64) | 0.5 |
| cg16171771 (PDCD6IP) | C-reactive protein | 7.02(4.63) | 0.13 | 23.46 (11.42 ) | 0.04 |
| cg16171771 (PDCD6IP) | WHR | -0.3(0.14) | 0.03 | -0.01(0.09) | 0.85 |
| cg01178710(MIR1287) | C-reactive protein | 11.04(4.66) | 0.02 | 29.3 (11.95 ) | 0.02 |
| cg01178710(MIR1287) | Creatinine | -0.11(0.29) | 0.7 | 3.64 (1.52 ) | 0.02 |
| cg01178710(MIR1287) | Uric acid | -0.93(2.43) | 0.7 | 6.06 (2.78 ) | 0.03 |
| cg04673737(5s_rRNA) | C-reactive protein | 10.18(3.39) | 2.9x10<sup>-3</sup> | 26.76 (8.78 ) | 0.003 |
| cg04673737(5s_rRNA) | Creatinine | -0.41(0.21) | 0.05 | 2.84 (1.12 ) | 0.01 |
| cg04673737(5s_rRNA) | WHR | -0.10(0.11) | 0.32 | 0.17 (0.07 ) | 0.02 |
| cg04673737(5s_rRNA) | BMI | 12.7(4.94) | 0.01 | 1.34(5.44) | 0.8 |
| cg24458314 (HDAC9) | C-reactive protein | 8.55(4.05) | 0.04 | 23.46 (10.39 ) | 0.03 |
| cg11024682(SREBF1) | Fasting glucose | -34.62(17.85) | 0.05 | 167.85(103.48) | 0.11 |
| cg11024682(SREBF1) | Insulin | 23.11(11.65) | 0.05 | -46.59(53.47) | 0.38 |
| cg11024682(SREBF1) | HbA1c | -0.19(0.72) | 0.79 | 7.42 (3.59 ) | 0.04 |
| cg11024682(SREBF1) | Triglycerides | 271.72(112) | 0.02 | 571.88 (219.02 ) | 0.01 |
| cg11024682(SREBF1) | BMI | 15.74(6.65) | 0.02 | 8.10(7.15) | 0.25 |
| cg11024682(SREBF1) | Creatinine | 0.58(0.29) | 0.04 | 1.99(1.50) | 0.19 |
| cg23372795(KCNK16) | LDL | -89.92(38.21) | 0.02 | 10.49(44.07) | 0.81 |
| cg23372795(KCNK16) | Cholesterol | -98.93(44.77) | 0.02 | -32.01(55.24) | 0.56 |
| cg23372795(KCNK16) | C-reactive protein | 9.56(3.31) | 0.004 | 26.35 (7.93 ) | 1.03x10<sup>-3</sup> |
| cg23372795(KCNK16) | Creatinine | -0.24(0.21) | 0.24 | 2.64 (1.02 ) | 0.01 |
| cg23372795(KCNK16) | Waist-hip ratio | -0.21(0.1) | 0.04 | 0.13 (0.06 ) | 0.05 |
| cg00574958(CPT1A) | Insulin | -31.25(15.25) | 0.04 | -102.46(72.94) | 0.16 |
| cg00574958(CPT1A) | Triglycerides | -426.52(145.67) | $3.7 \times 10^{-3}$ | -862.57 (298.33 ) | $4 \times 10^{-3}$ |
| cg00574958(CPT1A) | C-reactive protein | -22.03(5.96) | $3 \times 10^{-4}$ | -33.10 (15.94 ) | 0.04 |
| cg00574958(CPT1A) | WHR | -0.46(0.19) | 0.01 | -0.06(0.12) | 0.63 |
| cg00574958(CPT1A) | Uric acid | -9.73(3.08) | $1.8 \times 10^{-3}$ | -8.78 (3.69 ) | 0.02 |
| cg15815318 (RTN1) | C-reactive protein | 11.95(3.53) | $8 \times 10^{-4}$ | 22.43 (9.29 ) | 0.02 |
| cg15815318 (RTN1) | WHR | -0.13(0.11) | 0.21 | 0.17 (0.07 ) | 0.02 |
| cg15815318 (RTN1) | BMI | 10.23(5.16) | 0.04 | -7.19(5.62) | 0.2 |

*The effect size and p-value are shown in the table obtained after linear regression considering the risk factors as the dependent variable and DNA methylation of the CpG sites as the independent variable in the discovery phase sample of the study. A p-value of 0.05 has been considered significant. SE: Standard error, BMI: Body Mass Index; WHR: Waist hip ratio; LDL: Low-density lipids.*

### Identified signals are co-methylated in T2DM patients and associated with its risk factors

Since methylation at a single CpG site usually has a smaller effect on the disease risk, many CpG sites in the same gene or in a group of genes may work in tandem in a coordinated fashion to confer risk to the disease. To identify such co-methylated CpG sites, we constructed a methylation-based network of 535 differentially methylated CpG sites in 163 controls from the discovery phase of the study using WGCNA package in R. Our analysis identified 4 distinct modules, namely turquoise (296 CpG sites), blue (123 CpG sites), brown (28 CpG sites), and yellow (21 CpG sites), as shown in Figure 2a. We validated the preservation of modules in 254 independent T2DM subjects (Figure 2b) and in another data set of 55 independent individuals (28 individuals of Northern European ethnicity living in the USA for several generations and 27 migrant Indians to the USA who have migrated for at least five years) (data not shown). All 4 modules showed moderate to strong preservation of co-methylation pattern in both the validation datasets (Zsummary >2, Figure 2b) suggesting that co-methylation patterns between these CpG sites are reproducible across independent sample sets. Enrichment analysis for nearby genes of module’s CpG sites revealed significant enrichment of purine metabolism pathway genes (e.g., *ENTPD4*, *ADK*, *POLR3D*, and *POLE* genes) in the blue module and 8 other pathways (e.g., wnt signalling pathways, calcium signalling pathways) in the turquoise module (Supplementary Table 3).

#### MethQTL in JUP gene regulates hub genes of brown module

We further explored the brown module in detail as its eigengene was associated with the maximum number of T2DM risk factors (BMI, C-peptide, insulin, and systolic as well as diastolic pressure, Figure 2c). Since the structural properties of a network module can reveal its possible function and identify the hub genes regulating the modules, we checked the structure of the brown module and explored the connection pattern between the member CpG genes. Connectivity analysis of the module revealed 14 CpG sites as hub (connectivity > 0.8) members (Supplementary Table 4). With the help of Cytoscape, we visualized the module which revealed 2 submodules connected by CpG in *JUP* gene (Supplementary Figure 4). This suggests that methylation of CpG in *JUP* may regulate other genes in the brown module and different submodules may be regulated by independent mechanisms.

To find the possible regulatory mechanism of hub genes of the brown module, we performed methQTL analysis for SNPs of *JUP* gene in Illumina 610K quad BeadChip and methylation level of hub 14 hub CpG sites. Association analysis revealed that SNP rs6503650 in *JUP* gene significantly associates with methylation level of all 14 CpG sites that have been identified as hub CpG sites for brown module including cg15223933 in *JUP* gene itself (Table 3). Results from methQTL analysis and structure of brown module correlated with each other, indicating that the *JUP* gene and its variant rs6503650 plays important role in regulating module methylation behavior. Furthermore, it indicates that methylation of brown module genes is regulated by a common mechanism to mark their effect.

**Table 3:** Association of rs6503650 with methylation level of hub genes in brown module in methQTL analysis in 250 control subjects

| S. No. | CpG | Gene | SNP | Beta | SE | P-value |
| --- | --- | --- | --- | --- | --- | --- |
| 1 | cg06055730 | *FAM54A* | rs6503650(C/A) | 0.013 | 0.004 | 3.1x10<sup>-04</sup> |
| 2 | cg04583163 | *PIWIL3* |  | 0.009 | 0.003 | 2.03x10<sup>-03</sup> |
| 3 | cg13675624 | *PLEKHG6* |  | 0.01 | 0.003 | 2.40x10<sup>-03</sup> |
| 4 | cg17012502 | *ACVR1C* |  | 0.008 | 0.003 | 3.23x10<sup>-03</sup> |
| 5 | cg15223933 | *JUP* |  | 0.01 | 0.004 | 3.74x10<sup>-03</sup> |
| 6 | cg02854490 | *NOBOX* |  | 0.01 | 0.003 | 4.11x10<sup>-03</sup> |
| 7 | cg13936208 | *RNF34* |  | 0.008 | 0.003 | 5.50x10<sup>-03</sup> |
| 8 | cg08565320 | *TRNA Asp* |  | 0.009 | 0.003 | 5.85x10<sup>-03</sup> |
| 9 | cg07067773 | *ABRA* |  | 0.01 | 0.004 | 6.08 x10<sup>-03</sup> |
| 10 | cg18117601 | *ZSCAN18* |  | 0.008 | 0.003 | 0.01 |
| 11 | cg09510269 | *HCG18* |  | 0.008 | 0.003 | 0.02 |
| 12 | cg10620911 | *AGPAT1* |  | 0.009 | 0.004 | 0.02 |
| 13 | cg26020008 | *SPATA2* |  | 0.008 | 0.003 | 0.02 |
| 14 | cg11021810 | *FOXN3* |  | 0.01 | 0.005 | 0.05 |

The association status of rs6503650 (*JUP*) with methylation status of 14 hub CpGs in brown modules. A1/A2 = Allele1 and Allele2 for SNPs; Beta= Effect of in per A1 allele of SNP on methylation level of CpG sites. P-value =P-value of association for T2DM. The p-values, beta values, and standard errors have been obtained using linier regression considering the methylation at the CpGs as dependent variable and genotype at rs6503650 as independent variable.

#### Hub genes of the brown module segregate T2DM patients with dyslipidemia with good glycemic control

We performed hierarchical clustering of the T2DM patients using the DNA methylation level of 14 CpG sites. The combined power of these CpG sites stratified the T2DM cases into 3 groups with low, intermediate, and high levels of methylation CpG sites. Next, a comparison of the clinical parameters between these case groups showed significantly higher-level of triglycerides (P=0.01), lower level of HDL (P=0.005), and HbA1c (P=0.03) in patients with low methylation case group as compared to high methylation group was made This segregation of patients with distinct clinical phenotypes suggests that methylation at the hub CpG sites of the brown module can be used as useful markers to segregate patients with good glycemic control but with higher lipid level in clinics.

## Discussion

T2DM has multiple risk factors (e.g., high-fat diet, genetic susceptibility), and many of them are prevalent at the population level (4,5,6,7). Several of these factors can affect the disease risk through hypo- or hypermethylation of genes leading to alteration in specific gene expression (11,12,13,18). To identify these differentially methylated CpG sites, we performed an EWAS in 844 Indian individuals of Indo-European origin and explored the possible mechanism of associations by integrating epigenome-based co-methylation network analysis and methQTL analysis. To the best of our knowledge, this is the first study that integrates EWAS with a co-methylation study to identify DNA methylations that confer T2DM risk.

The current study identified a significant association of methylation in 3 known and 6 novel differentially methylated CpG sites with T2DM risk in Indian populations. We replicated hypomethylation of cg19693031 (in 3’UTR region of *TXNIP*) in T2DM cases like that of the non-resident Indian individuals in the London Life Sciences Prospective Population cohort (LOLIPOP) study (44). The association of cg19693031 methylation with T2DM risk has been replicated in the German (46), Caucasian (47), and Arab populations (48). As previously reported in Caucasian patients recruited in Spain, cg19693031 was also associated with sustained hyperglycemia (HbA1c) level in the Indian population (44). *TXNIP* is induced by higher glucose levels in pancreatic β-cell (49) and has been implicated in glucose-mediated apoptosis of β-cell death (50) in diabetes cases.

We also replicated hypermethylation of cg11024682 (*SREBF1*) as a risk factor for T2DM that was associated with incident diabetes in non-resident Indians in LOLIPOP cohort (44) and in the EPIC-Norfolk cohort (47). However, this signal failed to replicate in the Caucasian Botnia Cohort study (51) indicating a population-specific role towards T2DM risk. *SREBF1*, a transcription factor activated by AKT/mTOR signaling is responsible for the transcription of fatty acid synthesis genes (52) indicating that it may affect T2DM etiology through lipid-metabolism-mediated pathways. We also replicated the association of cg00574958 in *CPT1A* in the Indian population. Cg00574958 associated with incident T2DM in The Framingham Heart Study, EPIC-Norfolk, and LOLIPOP cohort (47). Further methylation of cg00574958 has also found to be associated with T2DM in Qatari family study (48), with adiposity among Ghanaians (53), with cardiovascular risk in *Registre Gironí del Cor* (Girona Heart Registry) of Spanish population, as well as with hypertriglyceridemia and waist phenotype of Mexican family study (54). *CPT1A* encodes for carnitine palmitoyltransferase 1 that transports long-chain fatty acids into the mitochondria for oxidation and contributes to maintaining glucagon secretion from pancreatic islets (55). Further, prolonged inhibition of *CPT1A* leads to lipid accumulation and insulin resistance in muscle cells of rats (56).

We also identified cg16171771 (near *PDCD6IP*), cg16171771 (near *MIR1287*), and cg04673737 (near *5S_rRNA*) as EWAS signal for T2DM (1.03×10^-11^) and are hypomethylated in the T2DM cases. Programmed cell death 6 interacting protein (PDCD6IP), is one of the most intensely studied multifunctional cytosolic and multi-domain scaffold proteins involved in multiple function including clathrin-mediated endocytosis, apoptosis, and also perform membrane repair (57). The gene may be involved in insulin secretion from pancreas via exocytosis. *MIR1287* is a microRNA that targets RAF1 which is involved in RAS-RAF-MEK-ERK pathways (58); hence, it may influence multiple metabolic process (e.g., glycolysis) relevant to T2DM physiology. *MIR1287* was downregulated in symptomatic patients with carotid plaques undergoing carotid endarterectomy (59) suggesting its possible role in metabolic disorders. Identified gene 5S_rRNA is a ribosomal RNA found in mitochondria and believed to play an important role in assemblage of 5 S subunit of RNA. It has been also suggested to have role in translational efficiency of mitochondrial ribosome and has differential modifications in T2DM mouse model as compared to healthy control (60). However, the precise role of these genes in T2DM physiology is largely unknown.

Our study identified hypomethylation at the gene-body CpG (cg24458314) in *HDAC9* as EWAS signal for T2DM. *HDAC9* encodes for the histone deacetylase 9, and the knockdown of *HDAC9* protects from adipose tissue dysfunction and improves glucose tolerance as well as insulin sensitivity in high-fat-fed mice (61). However, the exact relation between cg24458314 methylation and its role in *HDAC9* expression in T2DM needs to be explored for further mechanistic insight. Inhibition of this gene induces insulin resistance and T2DM in the case of high diet-fed mouse (61,62). We also identified cg15815318 in the *RTN1* gene body as hypomethylated in diabetic patients. *RTN1* codes for Reticulon 1 that is an established marker of chronic kidney disease and is highly expressed (63) in the kidneys of both human and murine models of HIV-associated nephropathy (HIVAN) as well as diabetic nephropathy. *In vitro*, RTN1A causes hyperglycemia and albumin-induced ER stress and death of both podocytes and renal tubular cells indicating that RTN1 is a key mediator for proteinuria-induced tubular cell toxicity and renal fibrosis in case of diabetic nephropathy (64). However, these signals need replication in other ethnic cohorts.

Additionally, we also identified cg23372795 site in *KCNK16* (potassium two pore domain channel subfamily K member 16) gene, a known GWAS signal for T2DM involved in insulin secretion in pancreatic β-cell as differentially methylated at EWA significance level. *KCNK16* induces β-cell membrane depolarization by increasing Ca2+ influx and tunes glucose-stimulated insulin secretion (65).

The comethylation network analysis in 264 controls revealed the presence of 4 methylated modules (turquoise, blue, brown, and yellow) among differentially methylated CpG sites that were preserved in independent datasets containing 254 T2DM cases suggesting a conserved co-methylation interaction among this CpG sites across samples. Further, the group methylations (eigengene) of the brown and yellow modules were associated with T2DM associated risk traits (e.g., BMI, CpG sites were correlated with T2DM risk factors). Detailed analysis of brown module identified 14 hub CpG low methylation sites that can be used to identify a subgroup of T2DM patients with significantly low HDL, high triglycerides, high C-peptide, but with low HbA1c. These patient groups are dyslipidemic with good glycemic control, which is uncommon to the usual case of dyslipidemia and poor glycemic control observed in most of the T2DM cohorts. Methylation at these 14 CpG sites can be used as prognostic markers to predict treatment outcome (e.g., lipid control, glycemic control) in patients, however these results need to be evaluated in a prospective cohort in the future.

Our analysis identified rs6503650 in the *JUP* gene as a potential methylation QTL for all 14 hub CpG sites of the brown module, indicating that *JUP* may regulate the methylation of other hub genes. *JUP* codes for junction plakoglobin and is required for insulin-induced signaling and glucose uptake in adipocytes (66). Plakoglobin binding promotes PI3K-Akt-FoxO signaling and improves glucose uptake and insulin sensitivity in muscles (67). The eigengene of the brown module was associated with insulin and C-peptide level along with BMI. The identification of the *JUP* gene as a central regulator for the brown module and its role in glucose uptake and insulin sensitivity suggests that the brown module genes may regulate glucose uptake and insulin sensitivity in muscle. However, this finding needs to be confirmed through functional studies in animal model systems and larger prospective human cohorts.

We recognize several limitations in our study. The discovery phase results showed a higher inflation factor that could be led to false-positive results in the discovery phase. However, replication of the signal in an independent validation cohort confirms the association of replicated signals. This co-methylation network analysis aimed at identifying the interactions between CpG sites needs validation through functional studies in appropriate model systems (e.g., cell lines).

In conclusion, the current study replicated 3 loci and identified 6 novel, robust and consistent CpG sites with differential methylations implicated for T2DM risk at epigenome-wide significance level. These appear to be related to T2DM via glucose and obesity-related pathways that had their effects in a coordinated manner as observed from the co-methylation analysis.

## Conflict of interest and author contribution

AKG researched data, performed the experiment contributed to discussion, and wrote the manuscript. GP, VK, and DK helped in the data collection. NT helped in study participant recruitment. DB researched data, contributed to discussion, and reviewed and edited the manuscript. DB is the guarantor of this work and, as such, had full access to all the data in the study and takes responsibility for the integrity of the data and the accuracy of the data analysis.The authors declare no conflict of interest.

## Acknowledgments

This study was supported by CARDIOMED (BSC0122-16) funded by the Council of Scientific and Industrial Research (CSIR), Government of India and University Potential of Excellence (UPOE II) from Jawaharlal Nehru University, New Delhi, India. The authors are thankful to all the participating subjects, their parents, and the school authorities for support and cooperation in carrying out the study. We thank Soham Bharadwaj for critically evaluating and editing the manuscript.

