## Supplementary material and additional file for "Network analysis after epigenome wide methylation study revealed JUP as a regulator of co-methylated risk-module for T2DM": Supplementary_file.pdf

### Supplementary Table 1: Sample characteristics of study participants

| | Discovery phase | | | Validation phase | | |
|---|---|---|---|---|---|---|
| **Trait** | **Cases** | **Controls** | **P-value** | **Cases** | **Controls** | **P-value** |
| Sex(M/F) | 135/125 | 134/130 | | 87/69 | 75/88 | |
| Age (years) | 46 (31-65) | 45 (40-79) | 0.30 | 54(47-60) | 50(43-60) | 0.03 |
| BMI ($kg/m^2$) | 24.34(22.49-27.12) | 24.34(22.07-28.08) | 0.18 | 26.14 (23.03-31.23) | 25.16(21.88-28.3) | 0.003 |
| WC (cm) | 86.36(81.28-91.44) | 89(81-95.75) | 0.19 | 91.44(86.36-100) | 87(80-94) | 0.002 |
| HC (cm) | 88.9(83.82-96.52) | 96(91-102.94) | $6.53x10^{-9}$ | 91.44 (85.75-99.06) | 97(91-104) | 0.002 |
| Waist-hip | 0.95(0.94-1.03) | 0.91(0.85-0.98) | $4x10^{-24}$ | 1.01 (0.96-1.05) | 0.90(0.84-0.96) | $3.47x10^{-17}$ |
| Fasting | 137.5(120-176.75) | 89.3(82.2-94.43) | $7.41x10^{-46}$ | 138 (110-175) | 89.15(83.32-96.23) | $1.72x10^{-32}$ |
| Fasting | 12.74(6.52-23.68) | 6.42(4.03-9.76) | $4.22x10^{-8}$ | 23.35 (11.44-36.85) | 7.22(4.2-11.7) | $9.07x10^{-20}$ |
| C-peptide | 3.19(1.85-4.69) | 1.79(1.26-2.56) | $2.17x10^{-12}$ | 3.66 (1.99-5.01) | 1.68(1.20-2.65) | $3.73x10^{-7}$ |
| HbA1C (%) | 8.35(6.72-9.82) | 5.3(4.97-5.59) | $2.15x10^{-60}$ | 7.8 (6.5-9) | 5.48(5.1-5.7) | $4.33x10^{-36}$ |

*Data has been represented as Median (Inter Quartile Range). P-value has been calculated using Wilcox test.*

**Supplementary Table 2**: List of CpGs selected for the validation

| CpG | Gene | Reason |
|---|---|---|
| cg19693031 | TXNIP | P-value |
| cg01178710 | MIR1287 | P-value |
| cg04673737 | 5S_rRNA | P-value |
| cg16171771 | PDCD6IP | P-value |
| cg16464924 | GAA | High Beta difference |
| cg00574958 | CPT1A | Associated with risk |
| cg06519434 | SCN5A | Associated with risk |
| cg07661849 | ZNF827 | Associated with risk |
| cg08679807 | ACOT7 | Associated with risk |
| cg08823985 | IL20RA | Associated with risk |
| cg11024682 | SREBF1 | Associated with risk |
| cg15815318 | RTN1 | Associated with risk |
| cg16934969 | RAPGEF4 | Associated with risk |
| cg16947583 | GULP1 | Associated with risk |
| cg23372795 | KCNK16 | Associated with risk |
| cg24458314 | HDAC9 | Associated with risk |
| cg26790091 | MAPK11 | Associated with risk |

**Supplementary Table 3:** Pathway enriched for genes corresponding to CpG sites present in identified two modules

| S.N. | Module | Pathways | Hyp_c | Genes |
|---|---|---|---|---|
| 1 | Turquoise | Arrhythmogenic right ventricular | $1.1x10^{-3}$ | *TCF7L1, CACNA1C, SLC8A1, ITGB6, CTNNA2, GJA1* |
| 2 | | Axon guidance | 0.01 | *ABLIM3, EPHB1, NFATC4, SEMA5B,* |
| 3 | | Wnt signaling | 0.01 | *WNT9A, LRP5, TCF7L1, NFATC4,* |
| 4 | | Basal cell carcinoma | 0.02 | *WNT9A, TCF7L1, FZD4, GLI2* |
| 5 | | Calcium signaling | 0.03 | *CACNA1C, CHRM2, SLC8A1, ATP2B2,* |
| 6 | | Hypertrophic | 0.04 | *CACNA1C, SLC8A1, ITGB6, PRKAG2* |
| 7 | | Protein digestion | 0.04 | *COL4A2, SLC8A1, ELN, SLC8A3* |
| 8 | | ECM-receptor | 0.04 | *SDC4, COL4A2, ITGB6, TNXB* |
| 9 | Blue | Purine metabolism | 0.04 | *ENTPD4, ADK, POLR3D, POLE* |

*Hyp_c = Hypo geometric test corrected p-value; for pathway enrichment analysis genes corresponding to respective CpG sites were entered in GENECODIS keeping human genome as background. Hyp_c p-value < 0.05 was considered significant for pathway enrichment analysis.*

**Supplementary Table 4:** Hub gene of brown module in the data

| S.No. | CpG | Connectivity | Gene |
|---|---|---|---|
| **1** | **cg10620911** | **0.91** | ***AGPAT1*** |
| **2** | **cg26020008** | **0.90** | ***SPATA2*** |
| **3** | **cg15223933** | **0.90** | ***JUP*** |
| **4** | **cg06055730** | **0.88** | ***FAM54A*** |
| **5** | **cg18117601** | **0.88** | ***ZSCAN18*** |
| **6** | **cg13675624** | **0.88** | ***PLEKHG6*** |
| **7** | **cg17012502** | **0.88** | ***ACVR1C*** |
| **8** | **cg02854490** | **0.88** | ***NOBOX*** |
| **9** | **cg13936208** | **0.88** | ***RNF34*** |
| **10** | **cg04583163** | **0.87** | ***PIWIL3*** |
| **11** | **cg07067773** | **0.86** | ***ABRA*** |
| **12** | **cg09510269** | **0.86** | ***HCG18*** |
| **13** | **cg08565320** | **0.83** | ***TRNA_Asp*** |
| **14** | **cg11021810** | **0.80** | ***FOXN3*** |
| 15 | cg01954686 | 0.78 | *ASB10* |
| 16 | cg17588904 | 0.71 | *HIPK1* |
| 17 | cg08872579 | 0.70 | *TTC22* |
| 18 | cg07661849 | 0.50 | *ZNF827* |
| 19 | cg07870129 | 0.39 | *KLF1* |
| 20 | cg03478313 | 0.17 | *RGMA* |
| 21 | cg23404610 | 0.08 | *FGFBP3* |
| 22 | cg25361454 | 0.02 | *ADAP1* |
| 23 | cg27266479 | -0.27 | *H6PD* |
| 24 | cg05491930 | -0.48 | *SFRS18* |
| 25 | cg01182973 | -0.55 | *ANKRD54* |
| 26 | cg11050859 | -0.59 | *TOX* |
| 27 | cg26187342 | -0.711 | *NOL11* |
| 28 | cg07624479 | -0.73 | *HTT* |

*Rows in bold are representing hub genes. Connectivity: Sum of all correlation values for all connected genes in topology overlap matrix.*

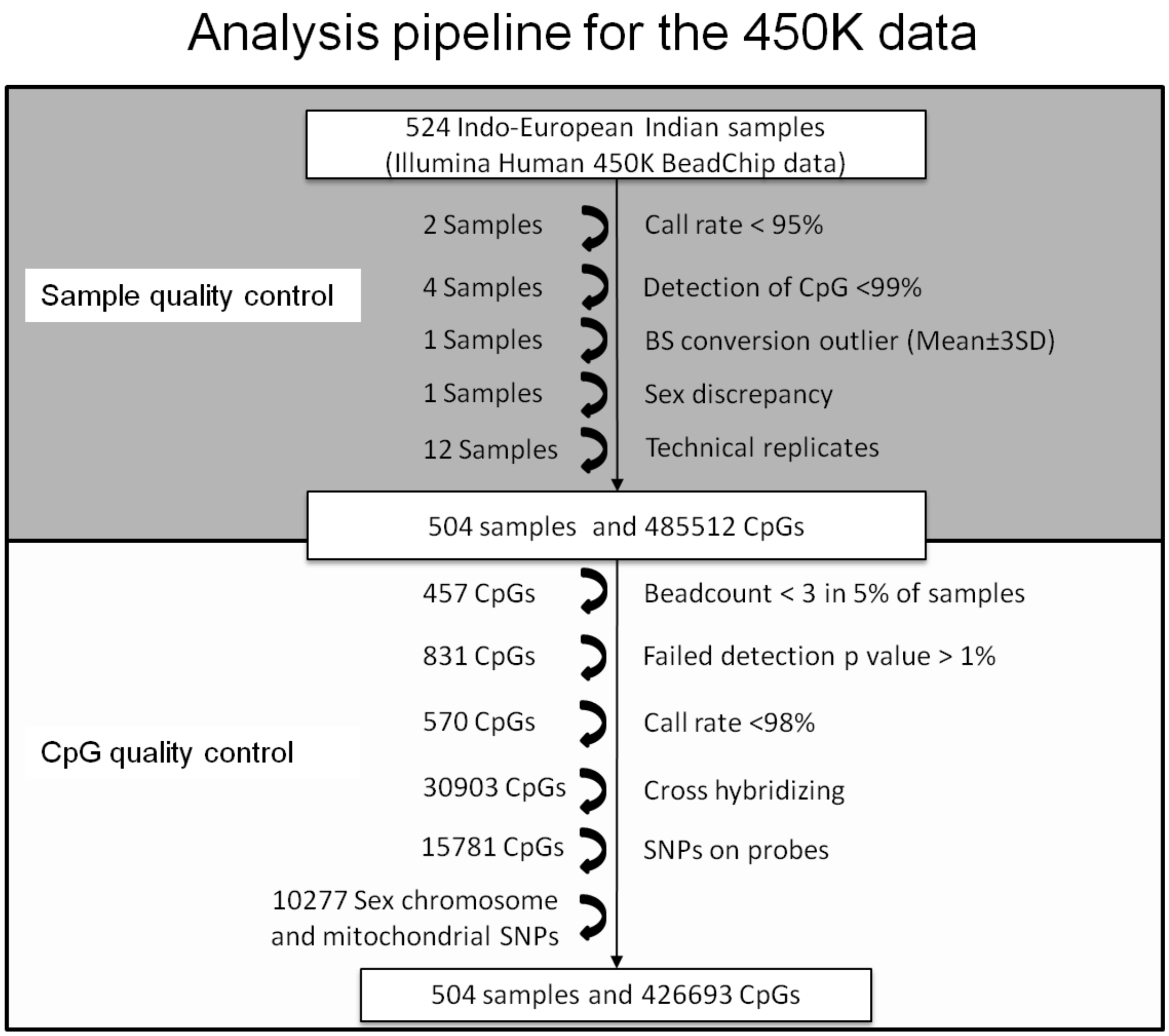


**Supplementary Figure 1:** *Detail of analysis pipeline (sample and CpG quality control steps) used in the study.*

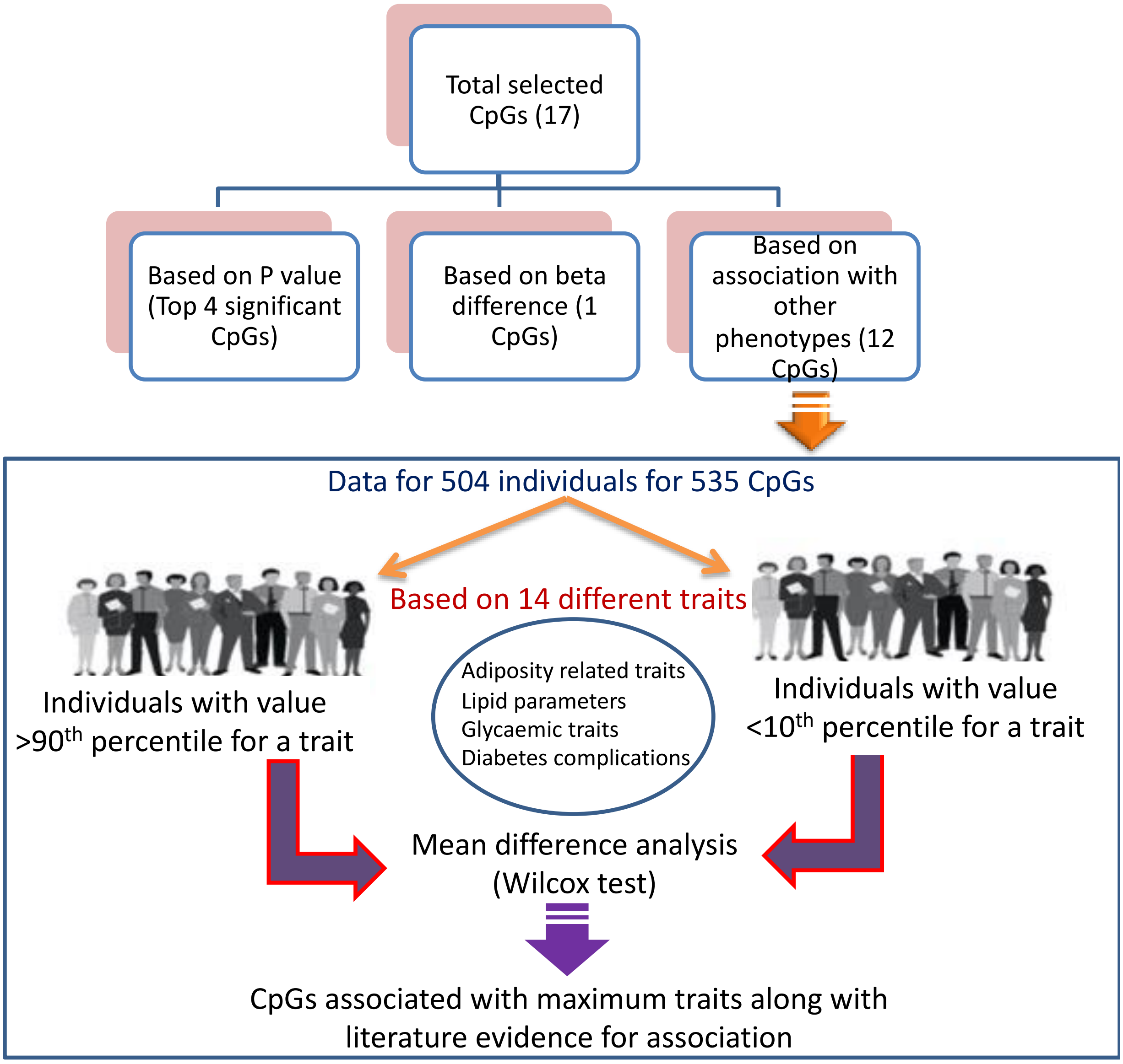


**Supplementary Figure 2:** *Detail of selection of CpGs for validation phase.*

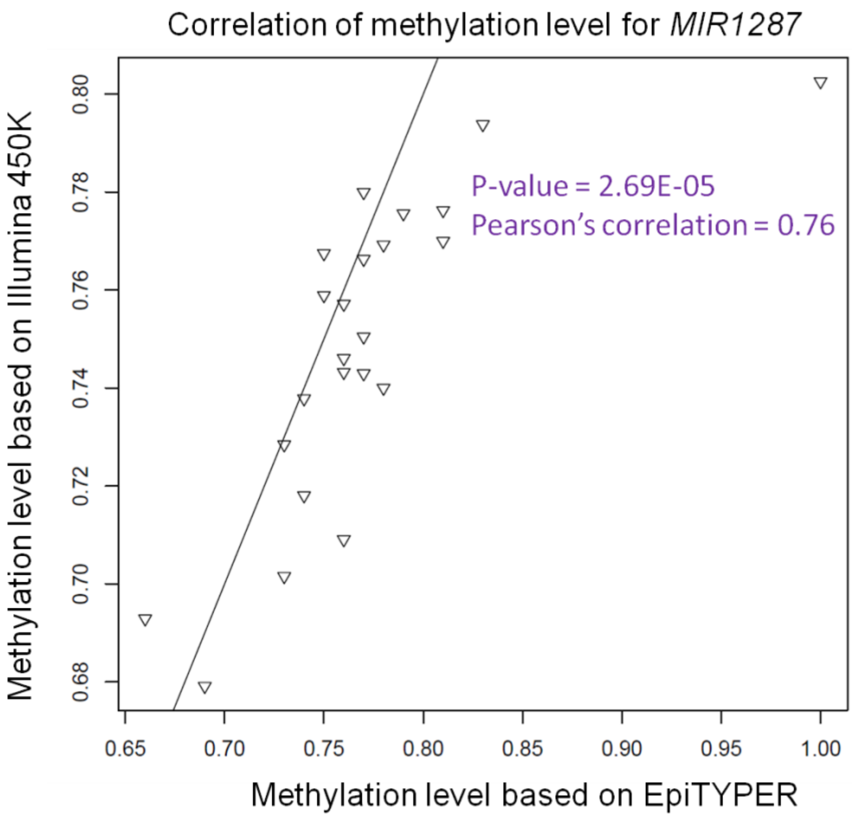


**Supplementary Figure 3:** *Scatter plot showing correlation between Illumina 450K and EpiTYPER data for cg01178710 in MIR1287 gene. The methylation values of cg01178710 for 25 samples have been generated using both Illumina 450K technology and EpiTYPER assay. Correlation has been calculated and scatter plot has been plotted using R. We have shown correlation for cg01178710 in MIR1287 as it was the best correlation observed between both technologies.*

a

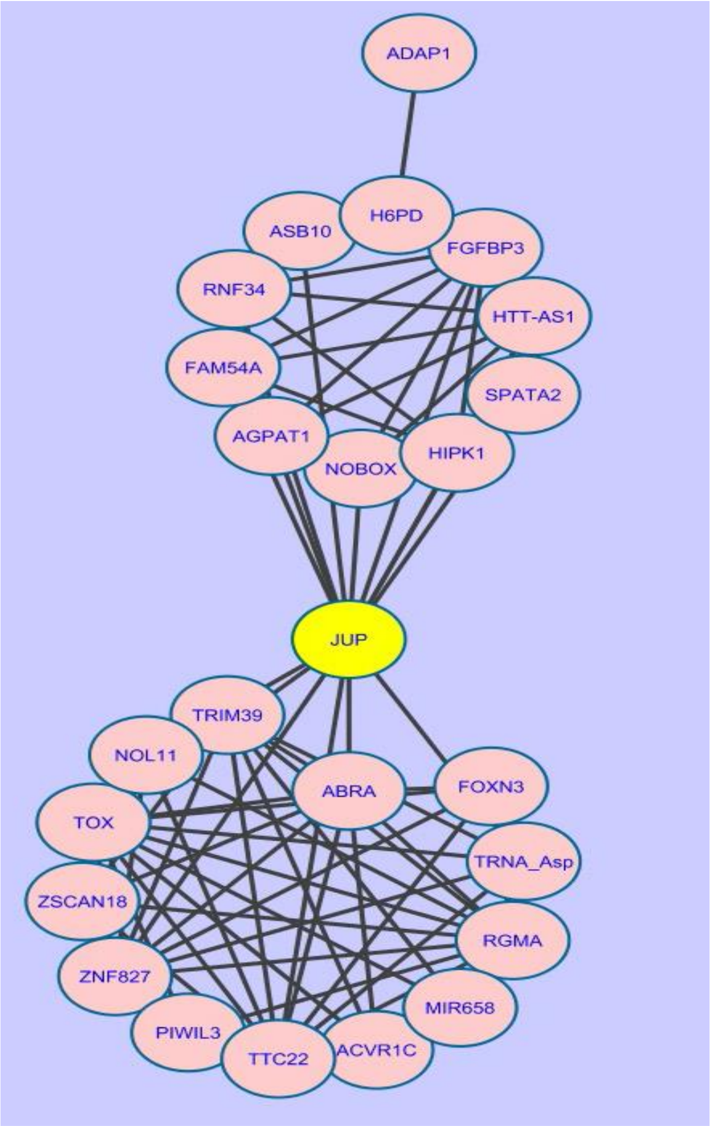


b

| **Node** | **Degree** |
|---|---|
| *JUP* | 15 |
| *TOX* | 13 |
| *RGMA* | 12 |
| *TTC22* | 12 |
| *ZNF827* | 11 |
| *TRIM39* | 10 |
| *ABRA* | 9 |
| *FGFBP3* | 9 |
| *HIPK1* | 8 |
| *HTT-AS1* | 7 |
| *TRNA_Asp* | 7 |
| *FAM54A* | 7 |
| *AGPAT1* | 7 |
| *RNF34* | 7 |
| *NOBOX* | 6 |
| *NOL11* | 6 |
| *ZSCAN18* | 6 |
| *FOXN3* | 6 |
| *ACVR1C* | 5 |
| *TOP1P2* | 4 |
| *ASB10* | 4 |
| *H6PD* | 3 |
| *MIR658* | 3 |
| *SPATA2* | 2 |
| *ADAP1* | 1 |

**Supplementary Figure 4:** *Visualization of brown module member genes in Cytoscape. The nodes with connectivity score >0.1 have been shown in the plot. (B) Degree of nodes (number of connections a node has to other nodes) in the brown module has been shown as Table.*
